# Motor improvement estimation and task adaptation for personalized robot-aided therapy: a feasibility study

**DOI:** 10.1101/728725

**Authors:** Christian Giang, Elvira Pirondini, Nawal Kinany, Camilla Pierella, Alessandro Panarese, Martina Coscia, Jenifer Miehlbradt, Cécile Magnin, Pierre Nicolo, Adrian Guggisberg, Silvestro Micera

**Author notes:** Equally contributing authors (Elvira Pirondini, Nawal Kinany, Camilla Pierella). **Corresponding author:** Christian Giang. **Co-authors emails:** Elvira Pirondini, Nawal Kinany, Camilla Pierella, Alessandro Panarese, Martina Coscia, Jenifer Miehlbradt, Cécile Magnin, Pierre Nicolo, Adrian Guggisberg, Silvestro Micera.

## Abstract

**Background:** In the past years, robotic systems have become increasingly popular in both upper and lower limb rehabilitation. Nevertheless, clinical studies have so far not been able to confirm superior efficacy of robotic therapy over conventional methods. The personalization of robot-aided therapy according to the patients’ individual motor deficits has been suggested as a pivotal step to improve the clinical outcome of such approaches.

**Methods:** Here, we present a model-based approach to personalize robot-aided rehabilitation therapy within training sessions. The proposed method combines the information from different motor performance measures recorded from the robot to continuously estimate patients’ motor improvement for a series of point-to-point reaching movements in different directions and comprises a personalization routine to automatically adapt the rehabilitation training. We engineered our approach using an upper limb exoskeleton and tested it with seventeen healthy subjects, who underwent a motor-adaptation paradigm, and two subacute stroke patients, exhibiting different degrees of motor impairment, who participated in a pilot test.

**Results:** The experiments illustrated the model’s capability to differentiate distinct motor improvement progressions among subjects and subtasks. The model suggested personalized training schedules based on motor improvement estimations for each movement in different directions. Patients’ motor performances were retained when training movements were reintroduced at a later stage.

**Conclusions:** Our results demonstrated the feasibility of the proposed model-based approach for the personalization of robot-aided rehabilitation therapy. The pilot test with two subacute stroke patients further supported our approach, while providing auspicious results for the applicability in clinical settings.

**Trial registration:** This study is registered in ClinicalTrials.gov (NCT02770300, registered 30 March 2016, https://clinicaltrials.gov/ct2/show/NCT02770300).

## 1 Background

With the increase of life expectancy, it is estimated that stroke related impairments will be ranked fourth most important cause of disability in western countries in 2030 [1]. With about 80% of stroke survivors experiencing significant motor impairment [2], stroke rehabilitation represents a major challenge. Despite early rehabilitative interventions, 55% to 75% of the patients still suffer from upper limb impairments in the chronic state of the injury [3–5]. The recovery of reaching and grasping movements is therefore a crucial therapeutic goal in stroke rehabilitation [6].

Post-stroke rehabilitation usually relies on task-oriented repetitive movements that help improving motor function and training new control strategies. In this regard, the amount of goal-directed and challenging practice, rather than daily intensity alone, seems to be the most effective factor in neurorehabilitation [7]. In the last two decades, robot-aided motor training has shown potential for the recovery of lost motor abilities in upper limbs after stroke [8–10]. While providing intense and highly repeatable motor training, robotic devices also offer means to control and quantify movement performances. Despite this undeniable potential, controlled clinical trials have so far not been able to confirm whether robotic therapy is more effective than conventional methods in restoring motor abilities [11, 12]. It has been argued that this might be related to saturation effects and a lack of automatic methods to promptly detect them [13].

The automatic and personalized adaptation of the rehabilitation training has been suggested as a pivotal step to improve the outcome of robot-aided rehabilitation and the clinical relevance of such solutions [14]. As a matter of fact, motor learning is known to be maximized when the difficulty level of the training task matches the patient’s level of ability [15]. Recent advances in the field of personalized robotic rehabilitation have therefore focused on the design of customized training protocols, including individualized selection of upper limb movements [16]. Different measures have been used to assess the patient’s “status” during training (i.e., motor performance, engagement, etc.) in order to adjust the proposed tasks accordingly. Kinematic performance measures, such as movement accuracy, smoothness, speed, inter-joint coordination, range of motion and stiffness [17–23], game-related statistics [13, 24], measures of muscle activity [17], or the combination of kinematic and psychophysiological measurements [25–27] have been among the measures used for the design of patient-tailored training protocols. However, those approaches either focused on a single performance measure describing a specific aspect of rehabilitation or used multiple measures but lacked the ability to meaningfully synthesize the information from all these variables. Integrating this information into a single measure, yet representative of the patient’s multidimensional rehabilitation response, would provide a straightforward method to track the multifaceted progress of the patient and trigger task adaptation while enormously simplifying the design of personalized rehabilitation training.

An interesting approach to address this issue was presented in the work of Panarese et al. [28]. The authors used a state-space model to merge the information from different kinematic measures and, in this way, estimated the motor improvement of chronic stroke patients exercising with a planar robotic device for upper limb rehabilitation. Similar methodologies have previously allowed to successfully characterize cognitive learning in animals [29, 30]. The results of Panarese et al. emphasized the potential of extending such approaches to the context of neurorehabilitation. In their study, the authors showed that the devised model was capable of mimicking decision rules applied by physical therapists regarding the adaptation of the task difficulty. In some cases, the model even appeared to be faster than the therapists in detecting when the patients’ motor performance had reached a plateau and when more challenging tasks should have been proposed.

In this work, we built on these results to implement a method able to continuously detect patient’s motor improvement and adapt the training task for three-dimensional movements using an upper limb exoskeleton. Indeed, most of the aforementioned adaptive approaches were restricted to planar workspaces, hindering their applications to functional movements exploring three-dimensional workspaces, which better resemble those performed during daily life activities. Evaluating and estimating motor improvement is particularly compelling in three-dimensional training workspaces, where the visual evaluation of motor performance becomes more challenging. Under these circumstances, a method able to autonomously estimate patient training progress, in particular for movements in different directions, could provide fundamental support to the therapists, enabling them to shift their focus from visual inspection of the movements performed to other important aspects of training. In this study, we also aimed at a continuous implementation of the motor improvement estimation and the personalization routine. Indeed, the immediate task adaptation within training sessions could not only increase patients’ engagement, but also foster their attention control, possibly leading to improved reaching performances [31].

In order to enable the use of such methods for clinical applications, it is first necessary to validate their feasibility and safety under controlled experimental conditions. We, therefore, devised an experiment to test our approach in a group of healthy subjects. In order to mimic the motor improvement observed in stroke patients, we applied a visual manipulation to the training environment. Previous studies on visually manipulated motor tasks showed that most people could cope with similar manipulations after training [32–36]. Accordingly, we hypothesized that performances would drop after the introduction of the inverted visual feedback (i.e., movements would become slower and less smooth), but would then gradually improve and eventually reach a plateau - with temporal dynamics resembling the ones occurring in robot-aided rehabilitation of stroke patients [28, 37, 38]. Using this setup, we tested whether our model was capable of tracking individual motor improvements induced by motor adaptation, and whether it was able to personalize the training by identifying “recovered” (i.e., adapted) movements in real-time. To provide further evidence about the feasibility and clinical usability of the presented approach, we finally performed a pilot test with two subacute stroke patients. The test was conducted in the framework of robot-aided upper-limb rehabilitation training for subacute stroke patients.

## 2 Results

### 2.1 Experimental validation with healthy participants

We first experimentally validated our model in a group of 17 healthy participants. Using the robotic upper limb exoskeleton ALEx [39, 40], we designed a three-dimensional point-to-point reaching task (Fig. 1a-b), a training exercise commonly used in robotic rehabilitation therapy [41–43]. In order to challenge the subjects and make them adapt to a new motor control scheme with temporal dynamics similar to those observed in robot-aided rehabilitation of stroke patients, the visual feedback was manipulated during five inversion blocks B_1-5_ (Fig. 1c, see Section 5.6.1). Under these circumstances, we tested whether our model was capable of continuously tracking MI (in this case induced by motor adaptation) and whether our implementation could personalize the training by identifying adapted movements (i.e., movements with performance comparable to the non-inverted condition) and by replacing them with more difficult ones.

**Figure 1.**
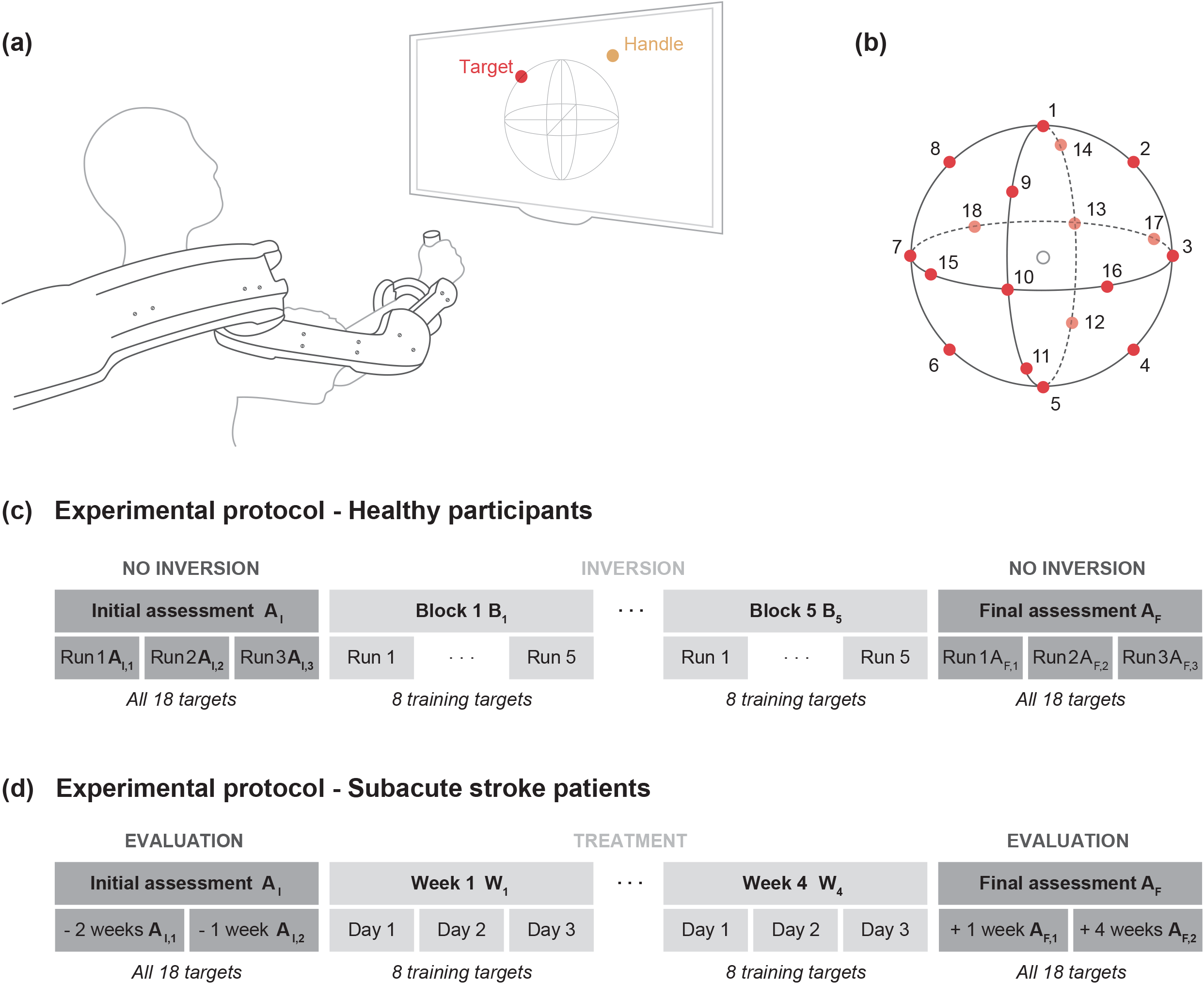
Experimental setup and protocols. (a) Schematic overview of experimental setup. (b) Design of the three-dimensional point-to-point reaching task. Eighteen targets (representing the different subtasks) are positioned over a sphere of 19 cm of radius (equally distributed on the three planes). The empty circle represents the center of the workspace (starting position). (c) Experimental protocol for healthy participants. Experiments were completed in a single session and were divided into blocks (one initial assessment block A_I,1-3_, five inversion blocks B_1-5_, one final assessment block A_F,1-3_). The assessment blocks consisted of three runs, each composed of 18 reaching movements (one towards each target). The inversion blocks consisted of five runs, each composed of eight reaching movements. The training targets for the inversion blocks were automatically selected by the implemented personalization routine. Breaks were allowed between the blocks to prevent fatigue. (d) Experimental protocol for the patient. During the initial (A_I,1-2_) and final (A_F,1-2_) assessment sessions, all eighteen targets were presented to the patient. For each treatment session eight training targets were selected by the implemented personalization routine. The total number of repetitions performed in each session was determined by the physical therapist.

#### 2.1.1 Task adaptation at subject level

Despite a general improvement for all participants, the subjects differed considerably in their adaptation speed, as quantified by the number of new targets introduced during the inversion blocks B_1-5_. We identified two groups using a median cut and found that the number of new targets for fast adapters (n = 9, 7.7±1.2 new targets, mean±std over subjects) and slow adapters (n = 8, 2.6±2.0 new targets) was significantly different (p < 0.001).

Interestingly, the two groups already showed differences in performance during the initial assessment A_I,1-3_ (Fig. 2a-c). Specifically, the values for MV were significantly higher (p < 0.001) for the slow adapters (0.171±0.041 m/s, mean±sem over subjects) in comparison to the fast adapters (0.159±0.040 m/s). In contrast, the values for SAL were significantly lower (p < 0.001) for the slow adapters (−3.134±0.715) compared to the fast adapters (−2.807±0.596). For the rate of SUCC no significant difference (p = 0.06) was found between slow (96.5±1.1 %) and fast (100±0.0 %) adapters.

**Figure 2.**
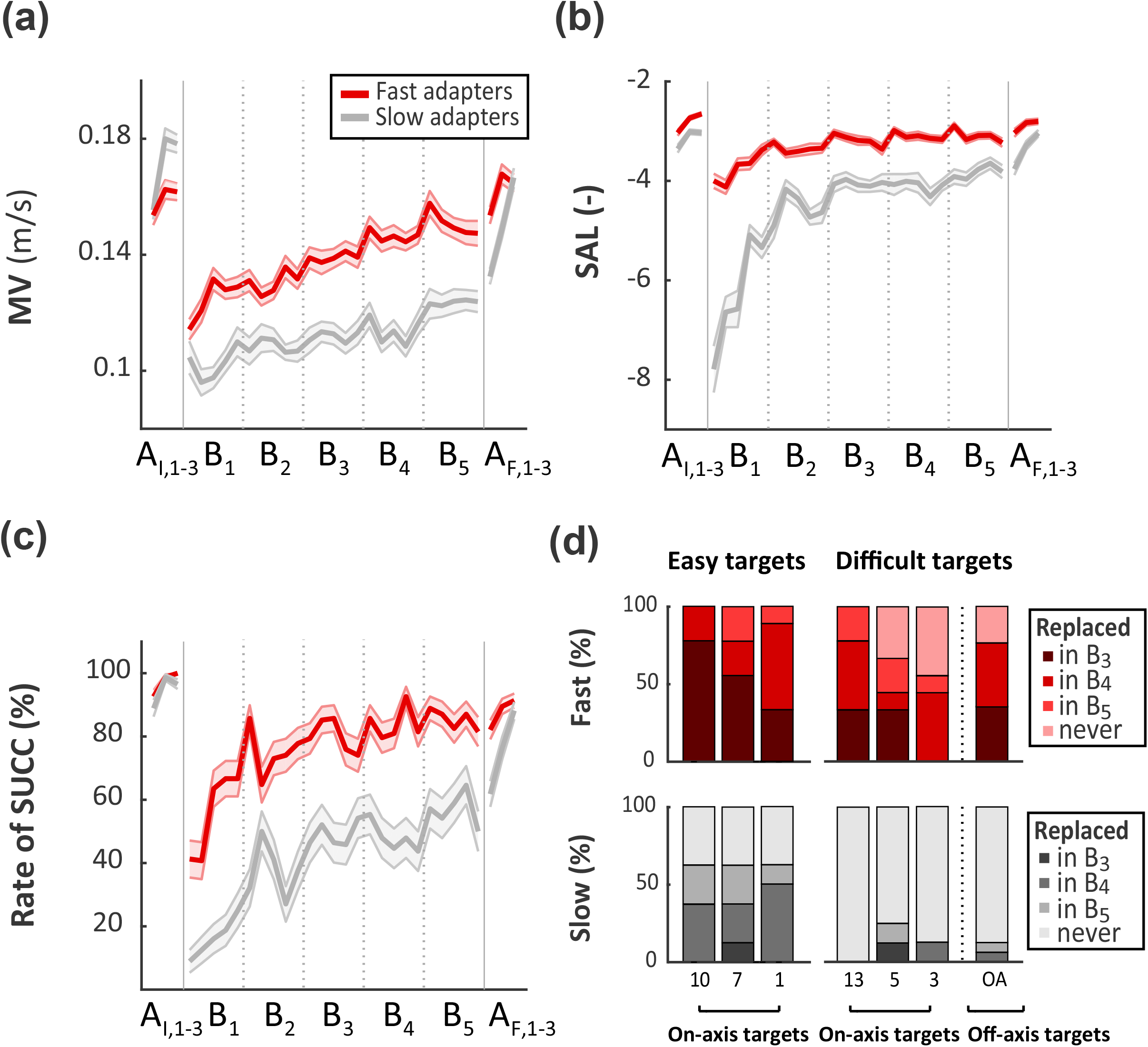
Analysis of performance measures for the experiment with healthy participants. Average values of mean velocity (MV, panel a), spectral arc length (SAL, panel b) and rate of SUCC (panel c) for each run (eight reaching movements) of fast (red) and slow (grey) adapters. Measures were averaged for all targets presented during a run and for all subjects of a group. Shaded areas depict standard error of the mean (sem). Vertical bars (panel d) depict the percentage of subjects in each group for which a target was replaced in B_3-5_ or was not replaced at all. No targets were replaced in and B_1-2_ due to lack of data needed for proper estimation of motor improvement.

As expected, the performance measures worsened for both groups after the introduction of the visual manipulation. However, the drop was remarkably smaller for the fast adapters: between the last run of the initial assessment A_I,3_ and the first run of the inverted block B_1_, the values for MV worsened by 0.048 m/s (−29% compared to A_I,3_) for the fast adapters and by 0.074 m/s (−41%) for slow adapters. The values for SAL worsened by 1.346 (−50%) for the fast adapters and by 4.802 (−157%) for the slow adapters. The rate of SUCC worsened by 57% for the fast adapters and 87% for the slow adapters. A two-way ANOVA illustrated the different impact of the introduced inversion on each group measured by MV (F1,421 = 10.62, p = 0.001), SAL (F_1,421_ = 112.31, p < 0.001) and rate of SUCC (F_1,421_ = 30.38, p < 0.001).

Both groups gradually improved from B_1_ to B_5_, although they did not reach their initial motor performances (i.e., performances during A_I,1-3_). A comparison between the last run of B_5_ and the last run of the initial assessment A_I,3_ showed that the fast adapters were more successful in restoring their initial performances: compared to their baseline level, MV was lowered by 0.014 m/s (−9% compared to A_I,3_) for the fast adapters and by 0.051 m/s (−30%) for slow adapters). The values for SAL were lowered by 0.581 (−22%) for the fast adapters and by 0.783 (−25%) for the slow adapters. The rate of SUCC was lowered by 18% for the fast adapters and by 45% for the slow adapters. During the entire experiment, the fast adapters outperformed the slow adapters and reached better final values for all performance measures, (+0.024 m/s for MV, +0.583 for SAL and +31% for rate of SUCC for the fast adapters at the last run of B_5_).

These results illustrate that the improvements induced by motor adaptation exhibited subject-specific dynamics, prompting the need for a model capable of differentiating between time courses of MI at subject level. We observed a coherency between the chosen performance measures and the adaptation speed quantified by the number of new training targets introduced. Fast adapters exhibited remarkably better performance compared to the slow adapters and they were thus introduced to considerably more training targets.

#### 2.1.2 Task adaptation at subtask level

In addition to the ability to differentiate MI for different subjects, we were interested in assessing whether the model was able to monitor MI at subtask level in a three-dimensional environment. Therefore, we evaluated which initial training targets were replaced by the algorithm during the inversion blocks and when this replacement occurred (Fig. 2d). The insertion of new targets did not start before B_3_, as in B_1-2_ the amount of data for each training target was not sufficient to obtain proper MI estimations (see section 5.1). As hypothesized in the experimental design, movements towards the off-axis targets (2, 4, 6, 8, 11, 14, 15, 16, 17 and 18, Fig. 1b) seemed to be more difficult: on average, the algorithm replaced these targets for 13% of the slow adapters and for 77% of the fast adapters. The on-axis targets (1, 3, 5, 7, 10 and 13), instead, were replaced for 38% of the slow adapters and 87% of the fast adapters. However, we also observed differences within the on-axis targets: on average, targets 3, 5 and 13 were replaced for 13% of the slow adapters and for 74% of the fast adapters, while the replacement for targets 1, 7 and 10 was achieved by 63% of the slow adapters and by 100% of the fast adapters. Following this analysis, we identified the subsets of easy (1, 7 and 10) and difficult (3, 5, 13 and off-axis) targets. Interestingly, the results suggested that despite the differences in the overall performance, the subsets of easy and difficult targets appeared to be similar for both groups. Nevertheless, we observed an earlier replacement of the easy targets for the fast adapters: 56% of the easy targets were replaced in B_3_ (4% for slow adapters), 33% were replaced in B_4_ (38% for slow adapters), and 11% were replaced in B_5_ (21% for slow adapters). In contrast, for the difficult targets, the fast adapters also needed more time to achieve a replacement (if they were replaced eventually): 26% of the difficult targets were replaced in B_3_ (3% for slow adapters), 35% were replaced in B_4_ (5% for slow adapters) and 14% were replaced in B_5_ (5% for slow adapters).

To illustrate the behavior of individual participants at subtask level, we present the data of one exemplary subject from each group for the movements towards the same two targets (Fig. 3). We selected one target from the subset of the easy (target 10) and one target from the subset of the difficult (target 13) targets. The examples illustrate the different adaptation rates observed between subjects and targets. For the easy target, the performance measures for the fast adapter quickly improved and approached a plateau. The slow adapter, instead, showed difficulties until the fourth repetition, reflected particularly by SAL and SUCC. Nonetheless, starting from the fifth repetition, he/she also managed to adapt the movements to the distorted visual feedback and finally reached the conditions for the target replacement at the twelfth repetition. The difficult target, instead, appeared to be more challenging for both subjects. For this target, the fast adapter showed an improvement in all performance measures only after the tenth repetition and finally reached the conditions for the target replacement after eighteen repetitions. In contrast, the slow adapter did not manage to satisfy the conditions for a replacement. Despite a trend of improvement, the motor performance was never sufficient to trigger a replacement of the target.

**Figure 3.**
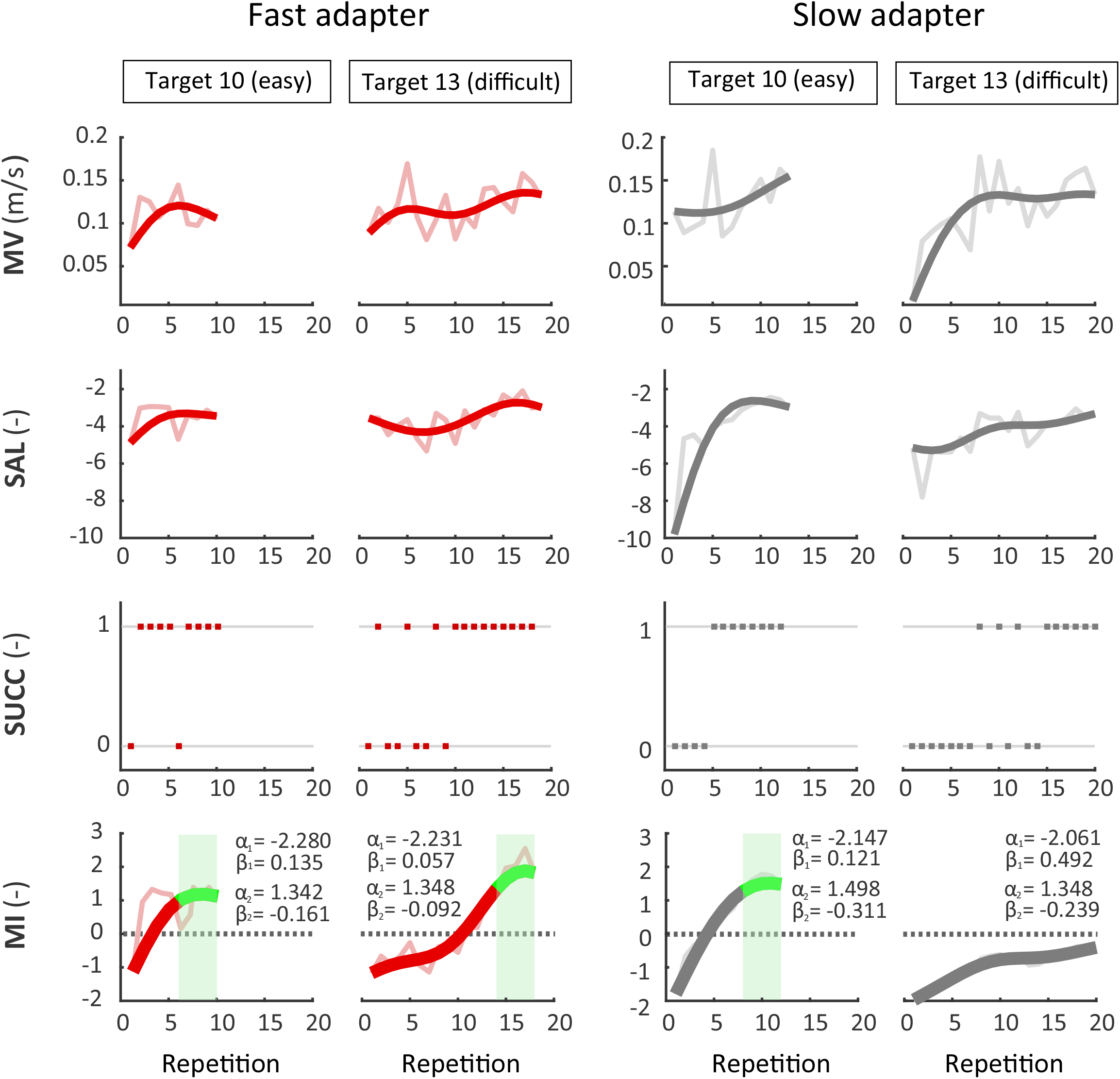
Examples of MI estimates and performance measures at subtask level. Data is presented for a fast adapter and a slow adapter for the same two targets. Repetitions for each target are concatenated for all inversion blocks and presented in chronological order. Data for mean velocity (MV), spectral arc length (SAL) and MI were low-pass filtered for visualization purposes (raw data shown in light red/grey). Dotted lines depict one of the necessary conditions (MI > 0) for triggering a target replacement. Green areas indicate the time span where the model detected a performance plateau and triggered a target replacement. Estimated model parameters (α_j_, β_j_) for each target and subject are presented next to the corresponding MI curves (a summary and analysis on the model parameters can be found in the *Supplementary material*).

These examples highlight the capability of the model to continuously capture individual time courses of improvement at subtask level. Furthermore, the results emphasized that the proposed model could detect saturation in motor performance and trigger actions regarding task difficulty in a well-timed manner.

### 2.2 Pilot test

To provide further evidence about the feasibility of the presented approach, we finally performed a pilot test with two subacute stroke patients, who completed four weeks of personalized robot-aided training in addition to standard rehabilitation therapy (Fig. 1d, see Section 5.5 for details). During the training, the set of targets was automatically adapted based on a continuous evaluation of the MI estimates for each training target.

Based on the initial assessment of their FMA-UE scores, we observed a remarkable difference in the degree of motor impairment of patient P01 (22 points at A_I,2_, Fig. 4a) compared to patient P02 (59 points at A_I,2_). This difference was reflected by the number of movements (nMov) performed in the training session, which was notably lower for P01 (31 movements compared to 69 movements for P02 at A_I,2_). The different degrees of initial impairment allowed us to evaluate the feasibility of our approach for two patients exhibiting disparate initial motor abilities.

**Figure 4.**
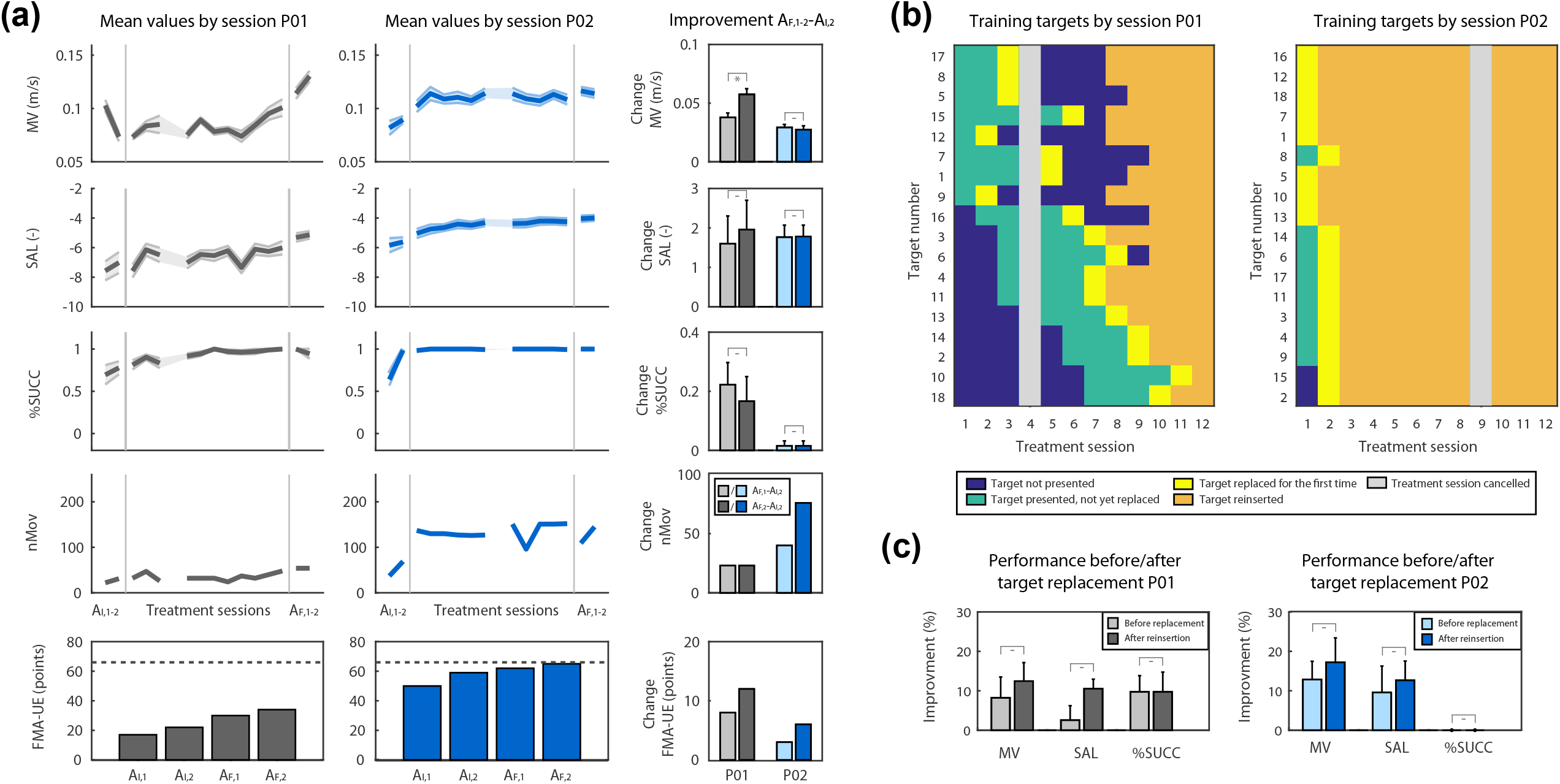
Summary of the results from the pilot test with two subacute stroke patients. (a) The first three rows show the mean values for mean velocity (MV), spectral arc length (SAL) and rate of success (SUCC) for each assessment and treatment session. Measures were averaged for all targets presented during a session, shaded areas depict standard error of the mean (SEM). The fourth row shows number of movements performed by the patients in each session. The fifth row shows the scores on the Fugl-Meyer scale for upper extremities (FMA-UE) for initial (A_I,1-2_) and final (A_F,1-2_) assessment sessions. The dotted line indicates the maximum achievable score for FMA-UE (66 points). Last column shows changes for all metrics between the final assessments A_F,1-2_ and the second initial assessment A_I,2_. Error bars depict standard error of the mean (SEM). Statistical significance between values are indicated by asterisks (*, p < 0.05) or dashes (−, p > 0.05). (b) Summary of the training targets presented to the patients in each treatment session. Targets are listed by the order as presented to the patients (first eight targets from the top are the initial training set). (c) Analysis of performance measures for two different time points (i.e., before replacement and after reinsertion). The data shows the mean values for MV, SAL and SUCC averaged for all targets at these time points. Values are compared between the last four movements towards a training target before its replacement and the first four movements towards the target after it has been reinserted for training. Values are given relatively to the mean values obtained from the first four movements towards all targets. Error bars depict standard error of the mean (SEM). Statistical significance between values are indicated by asterisks (*, p < 0.05) or dashes (−, p > 0.05).

Following the training, both patients showed improvements for MV, SAL, and SUCC. When comparing the values right before (A_I,2_) and right after (A_F,1_) the treatment sessions, we observed that patient P01 improved MV (+0.378 m/s), SAL (+1.60) and rate of SUCC (+22%). In comparison, improvements for patient P02 were lower for MV (+0.293 m/s), slightly higher for SAL (+1.77) and remarkably lower for SUCC (+1.6%). The latter can be explained by the fact that the values for SUCC for patient P02 already started at a very high level (98% at A_I,2_), leaving smaller room for improvement. Interestingly, both patients managed to retain or even improved their performance in the follow-up assessment (A_F,2_) four weeks after completion of the training. The only exception was observed for patient P01, who slightly worsened in SUCC between A_F,1_ and A_F,2_. However, this difference did not appear to be statistically significant (p = 0.61). Along with the improvements of the performance measures, we also observed higher FMA-UE scores for both patients following the training. In that respect, we also observed a lower increase for patient P02 (+3 points) compared to patient P01 (+8 points) between A_I,2_ and A_F,1_. Interestingly, both patients further improved their FMA-UE scores when assessed in the follow-up session A_F,2_. Finally, we also observed an increase in the number of performed movements per session (nMov) for both patients. As for this measurement, instead, patient P02 (+40 movements at A_F,1_ and +76 movements at A_F,2_ compared to A_I,2_) improved more than patient P01 (+23 movements at A_F,1_ and A_F,2_ compared to A_I,2_).

Both patients progressed during the rehabilitation training and eventually achieved a replacement of all eighteen training targets. However, the temporal dynamics of these replacements appeared to be strongly different for each patient (Fig. 4b). In line with the lower degree of motor impairments observed from the performance measures and the FMA-UE scores, patient P02 achieved a replacement of all training targets after only two training sessions. Patient P01, instead, needed considerably more time to achieve the replacement of all eighteen targets. While some of the initial training targets (i.e. targets 9 and 12) were already replaced after two treatment sessions, other targets (i.e. targets 1, 7 and 15) needed more than 4 training sessions to trigger a replacement. It was only after eleven treatment sessions that all eighteen training targets were presented to patient P01. These observations emphasized the ability of our model to differentiate between both subject- and subtask-specific time courses of motor improvement, also in a real clinical setting. The examples illustrate how the model adapted the training schedules according to the patients’ individual abilities, granting patient P01 enough time to practice the movements, and at the same time, responding to the fast recovery of patient P02 by continuously introducing new training targets.

Upon completion of the full set of training targets (i.e., when all targets had been replaced at least once), the therapy was carried on by reintroducing all targets and presenting them alternatingly in the order in which they were replaced. This allowed us assessing whether the patients’ performance was retained once a training target was reintroduced, so as to validate that the replacements orchestrated by the algorithm had occurred when the movements towards the targets had actually recovered. In order to do so, we compared the mean values for MV, SAL, and SUCC from the last four repetitions of a movement before a target was replaced by the algorithm with the mean values of the four repetitions of the same movement after the first reinsertion as a training target (Fig. 4c). Both values are presented relatively to the mean values obtained from the first initial four repetitions of the movements towards a training target. The overall analysis for all eighteen targets showed that compared to the initial movements towards the targets, almost all values for the three performance measures were higher (MV by +8% for P01 and by +12% for P02, SAL by +3% (+10%) and SUCC by +10% (+0%)) right before the targets were replaced by the algorithm. Moreover, both patients retained or even improved their performance for a movement when the corresponding training target was reintroduced at a later stage. For both patients, we found no significant difference (p > 0.077) for the values of all three performance measures between the two time points (i.e., before replacement and after reinsertion). These results illustrate that the algorithm only replaced training targets when motor performance had stably improved. More importantly, both patients have retained these improvements when the training targets were reintroduced at a later stage, suggesting that the timing of the replacement was appropriate.

## 3 Discussion

In this study, we presented and validated a model-based approach for the personalization of robotic rehabilitation training based on motor performance during three-dimensional training tasks. Capitalizing on the enhanced potential for plasticity in the early stage after the injury [44, 45], the model was designed to allow estimation of motor improvement (MI) in subacute stroke patients. A first experimental validation in healthy subjects demonstrated the ability of our model to capture MI linked to visual motor adaptation. The results were further validated by a clinical pilot test with two subacute stroke patients, in which motor recovery was tracked and harnessed by our personalization method.

### 3.1 Motor improvement model for 3D reaching tasks

One of the pivotal aspects underlying the development of a personalized rehabilitation training is the definition of performance measures that can correctly capture the different aspects of motor recovery, as well as their specific dynamics. Three performance measures were selected based on previous studies [28, 46] and used to devise a state-space model for MI estimation: movement velocity (MV), spectral arc length (SAL), and robot assistance dependency (SUCC). In past studies, the selected measures have been shown to correlate with clinical scores [47] and they have been linked to distinct post-stroke deficits and mechanisms of recovery [48, 49]. Specifically, the percentage of accomplished tasks was mostly associated to paresis (i.e., the decreased ability to volitionally modulate motor units activation [50]), whereas movement speed and smoothness were related to an abnormal muscle tone [48]. We therefore hypothesized that considering a combination of these measures was necessary to obtain a comprehensive assessment of the patient’s rehabilitative status. As such, we aimed to design a model capable of integrating the information coming from these multiple variables into a single motor performance measure, that could i) allow a better tracking of the patient’s rehabilitation progress, and ii) simplify the design of an automatic and personalized training protocol, therefore possibly enhancing the efficacy of the robot-aided rehabilitation training.

Using the robotic upper limb exoskeleton ALEx [39, 40], we designed a three-dimensional point-to-point reaching task, a training exercise commonly used in robotic rehabilitation therapy [41–43]. The movement amplitude was selected to allow the exploration of a functional workspace, while movement directions were chosen to elicit independent and synergistic motion of shoulder and elbow, capitalizing on the advantages provided by robotic exoskeleton devices [51]. Such design not only allowed the users to explore an extensive workspace, but also provided a way to easily assess their performance for the different regions of the workspace (i.e., for different subtasks, represented by the movements towards the different targets). The reaching task was displayed on a screen mounted in front of the participants and visual feedback was provided by means of a cursor mapping the position of the exoskeleton’s handle to the screen, an important aspect to avoid compensatory strategies [13]. The choice of a 2D screen was justified by the typically advanced age of post-stroke patients, who are usually not familiar and, therefore, often discomforted by 3D immersive reality. In order to preserve the depth perception, the dimension of the target spheres was modified in accordance with their position in the 3D space. Preliminary data from a group of age-matched healthy subjects (see *Supplementary*) showed that performance measures were not different for targets on the depth axes, confirming that the depth could be properly perceived by the users.

### 3.2 Adaptation to visually manipulated reaching tasks in 3D

We first sought to validate the model’s ability to continuously track MI and dynamically adjust the training task under controlled conditions. To this end, we presented a motor adaptation task to a group of seventeen healthy subjects. In order to mimic the motor deficits observed in stroke patients, we introduced a manipulation of the visual feedback, by inverting the directions of the 3D environment. While the physiological mechanisms underlying motor adaptation and motor recovery are most likely not equivalent, the main objective of this experimental design was merely to obtain an adaptation curve that resembles post-stroke motor recovery, on which we could validate the efficacy of our model. Our results indeed illustrated that motor adaptation in healthy subjects and motor recovery in stroke patients exhibited similar temporal dynamics.

During the experiments, the MI model tracked when a movement towards a target was performed efficiently despite the visual perturbation, and subsequently adjusted the training by replacing this target with a more difficult one from the training queue. Interestingly, we observed that the number of new training targets inserted strongly differed across participants, pointing out varying adaptation speeds. This result was not expected *a priori*, but it emerged as an unforeseen opportunity to highlight the model’s capability to differentiate individual motor adaptation rates. Based on the number of new inserted training targets, we divided the healthy population into two separate clusters: fast and slow adapters. The analysis on the performance measures showed that the fast adapters learned to cope with the manipulated environment very quickly, while the slow adapters needed considerably more time to reach similar performances. Interestingly, the two groups already showed differences in motor performances during the initial assessment. When the visual feedback was manipulated, the slow adapters presented a strongly reduced speed and motion smoothness. This was particularly the case earlier before the use of the adaptive algorithm and we, therefore, believe that the latter did not have an influence on the participants’ performance. The MI model, instead, was able to capture these individual performance differences at subtask level and coherently introduced new training subtasks in a well-timed manner, i.e., targets were replaced when subjects reached a performance plateau. The advantages of monitoring motor improvement at subtask level were supported by additional post-hoc analyses (see *Supplementary material*). The analyses illustrated that if motor improvements were estimated for the reaching task as a whole (i.e., combining the recorded data for movements in all directions), improvements for individual subtasks would have been obscured by inferior performances of other, more difficult, subtasks. Moreover, the detection of performance plateaus would not correspond to the actual performances for any subtask. As a result, some subtasks would be kept too long, while others would be replaced too soon, potentially leading to a less efficient training schedule. For instance, the overall MI estimate based on the first 80 repetitions of the slow adapter suggests a performance plateau already after 39 repetitions (which corresponds to approximately 5 repetitions for each subtask). However, when looking at the performance measures of this subject for target 13 separately, it is clear that a replacement of this target after 5 repetitions would have been premature. We therefore believe that this analysis further supports our approach to specifically consider MI estimation at subtask level.

As hypothesized in the experimental design, off-axis targets were replaced less often than on-axis targets and they, thus, seemed to be more difficult. However, the results showed that there were also remarkable performance differences among the on-axis targets. An analysis on the replaced training targets demonstrated that the subsets of easy (1, 7 and 10) and difficult (3, 5, 13 and off-axis) targets appeared to be similar for both types of adapters: easy targets were mostly replaced earlier and more frequently than the difficult ones. It could be that the medial and proximal movements towards targets 7 and 10 tended to be easier for the participants. However, since these tendencies were not observed in the patients or the healthy subjects involved in the preliminary study (see *Supplementary material*), we presume that the performance differences for the on-axis targets could be linked to the visually manipulated environment. Previous studies have investigated visual manipulation in planar reaching movements and suggested that the adaptation to such manipulations involves a complex mixture of implicit and cognitive processes [33, 52]. However, further research would be necessary to examine these phenomena in three-dimensional reaching movements. As a matter of fact, existing literature covering this area is still relatively sparse. In this context, it would be interesting to determine why the reaching movements towards some on-axis targets appeared to be more challenging in the inverted environment, independent from the individual adaptation speed of the subjects.

Finally, we would also like to raise the question of psychological implications resulting from the automated training adaption. From qualitative observations made during the experiments with the healthy subjects, we noticed that many participants showed increased motivation and verbalized satisfaction when new training targets were introduced. Motivation is known to be a crucial factor in rehabilitation and finding ways to maintain and improve it has always been a matter of interest [53–55]. With regard to this issue, it seems like the automated character of our approach, enabling dynamic and well-timed task adaptation, may have positive impacts on training engagement. Transferring this benefit to the rehabilitation program of patients may promote training motivation and hence potentially improve the clinical outcome of robot-aided rehabilitation trainings.

### 3.3 Personalization of rehabilitation therapy

The potential of our implementation was finally evaluated in a clinical pilot test with two subacute stroke patients, who completed four weeks of robot-aided rehabilitation training following our adaptive approach.

The results obtained from these two patients suggested that in general, the selected performance measures (MV, SAL and SUCC) appeared to be suitable for the use with the presented motor improvement model and the temporal dynamics appeared to be coherent with the chosen probability models and with results from previous work [49]. We observed improvements for all three performance measures following the training. Nevertheless, some tuning of the parameters could be considered to further enhance the efficacy of the motor improvement model. For instance, we observed that the patient with a lower degree of initial impairments (P02) barely made use of the robotic assistance provided by the exoskeleton, leading to almost no variance in the variable SUCC. In this regard, future studies may explore other performance measures and models, such as the ones proposed by Panarese et al. [28, 49], to achieve a more exhaustive evaluation of the patients’ status.

Based on the devised method, the training of the two patients following the personalized rehabilitation protocol was continuously monitored and the point-to-point reaching task was adapted in real-time to match their level of ability. The analysis showed that targets were indeed replaced by the model at appropriate moments, i.e., when the patients’ performance had improved and started to saturate. Indeed, it could be argued that a replacement of a subtask occurring too soon would have led to degraded motor performances in further evaluations. However, the results demonstrated that motor performances of both patients were retained when targets were reintroduced, indicating that the estimated recovery was preserved. Nevertheless, other methods for training scheduling could be introduced to further optimize the training progression. Indeed, previous work has suggested that effective scheduling of multitask motor learning should be based on prediction of long-term gains rather than on current performance changes [56]. Along these lines, we have implemented the time window of the last four repetitions, which are always taken into account for the evaluation of motor performance. However, it should be acknowledged that other, more sophisticated, methods to adapt the schedules may lead to higher gains in rehabilitation and are therefore worth exploring. For instance, task difficulty could be increased by introducing new subtasks depending on more complex movements within the same workspace, in order to exploit generalization effects [57, 58]. Another possible approach could be a semi-automatic implementation of the personalization, where the physical therapists remains in charge of the task adaptation, in order to benefit from their expertise, while in parallel harnessing the real-time MI estimates provided by the model as a decision support. Such solutions could further improve engagement and enhance the rehabilitative treatment by providing training tasks specifically adapted to the ability level of the patient.

As a way to measure the clinical outcome of the rehabilitative interventions, the patients completed the Fugl-Meyer assessment for upper extremities in all initial and final assessment sessions. When comparing the scores of the patients between the second initial assessment A_I,2_ and the first final assessment A_F,1_, we found that both patient P01 (+8 points) and patient P02 (+3 points) improved. Considering the scores at the follow-up session, A_F,2_, four weeks after completion of the training, this improvement was further sustained for both patients (+12 points for P01 and +6 points for P02). In addition to this gain in FMA-UE scores, the improvement of motor performance, along with its subsequent retention at target reinsertion, are promising indications for the usability and efficacy of the presented approach in clinical settings. Nevertheless, it is also well known that subacute patients often report motor improvements even with limited training [59]. Therefore, it cannot be presumed that improvements were merely elicited by the robotic rehabilitation trainings. However, several pieces of evidence suggested that the period immediately after the lesion, normally characterized by spontaneous neurological recovery, represents the critical time window in which the delivery of high dose and intense neurorehabilitation can elicit crucial improvements in functional tasks [60, 61]. Therefore, more and more robot-aided rehabilitation trainings should be targeting subacute stroke populations. In this context, our results illustrate the feasibility of using a personalization method to continuously monitor the status of both mild and severely impaired post-stroke patients and to automatically adapt their motor retraining within practice sessions. The latter might be particularly pivotal in the context of rehabilitation training for subacute stroke patients. In contrast to chronic patients, this population often shows potential for quick recovery [49], calling for prompt training adjustments in order to continuously challenge their neuromuscular system.

Nevertheless, further studies including larger cohorts of participants would be necessary to draw meaningful conclusions about the clinical relevance of the presented approach. In this context, it would be particularly interesting to compare the clinical outcomes of the personalized approach presented in this study with non-adaptive robotic or conventional rehabilitation trainings. Indeed, previous work has suggested that pseudo-random scheduling of multiple tasks may be almost as effective as adaptive scheduling approaches [56]. To demonstrate clinical relevance, it is therefore crucial to assess the efficacy of the presented approach in large clinical trials, focusing their activity on the comparison of adaptive and non-adaptive schedules. In this context, the results obtained from this work may provide a useful basis for the design and implementation of such clinical studies.

## 4 Conclusions

In this work, we presented a model-based approach to personalize robot-aided rehabilitation therapy within rehabilitation sessions. The feasibility of this approach was validated in experiments with seventeen healthy subjects and a pilot test with two subacute stroke patients providing promising results. However, due to the limited sample size, larger studies would be needed to demonstrate clinical relevance of the presented approach. While we implemented the proposed method for the use in upper limb rehabilitation of stroke patients, the usage is certainly not limited to such applications. The presented model can be adapted for the use with other robotic rehabilitation devices and training tasks, exploiting different performance measures and/or different observation equations. The real-time functionality and the identification of subject-specific abilities at subtask level could enhance robot-aided rehabilitation training, making it more purposive and efficient for the patients.

## 5 Methods

Based on the work of Panarese et al. [28], we developed a model to continuously estimate motor improvement (MI) in three-dimensional workspaces using kinematic performance measures. We then designed a personalization routine, which automatically adapts the difficulty of the rehabilitative motor task (i.e., a point-to-point reaching task) based on the MI estimates. Both the MI model and the personalization routine were integrated in the control algorithm of an upper-limb exoskeleton and tested with a group of 17 healthy participants. The presented approach was then tested with two subacute stroke patients.

### 5.1 Motor improvement model

In order to continuously track patients’ MI at subtask level (i.e., for a series of point-to-point reaching movements in different directions), we used a state-space model. MI was modelled as a random walk:

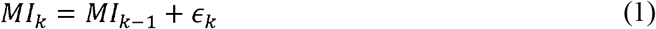

where *k* are the different repetitions for a movement direction and ϵ_k_ are independent Gaussian random variables with zero mean and variance 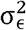. A set of observation equations z_j,k_ was defined in order to estimate MI. These equations related MI to continuous performance measures r_j_, which were computed from kinematic recordings provided by the robotic device (see section 5.2 for details on the performance measures). The continuous variables r_j_ (with *j* = 1,.., *J* representing the different performance measures) were defined by the log-linear probability model

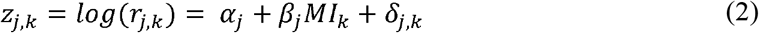

where δ_*j,k*_ are independent Gaussian random variables with zero mean and variance 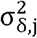. The use of log-linear models allowed capturing rapid increases (or decreases) of the performance measures during the training, as well as the expected convergence towards subject-specific upper (or lower) bounds at the end of the training. The suitability of such probability models for motor performance measures in stroke patients was previously demonstrated [28, 49]. Similarly, an observation equation for a discrete performance measure n_k_ was defined. The binary discrete variable n_k_ ϵ {0, 1} was used to track the completion of the exercised subtask, with 1 meaning that the subtask was performed successfully and 0 meaning failure. The observation model for n_k_ was assumed to be a Bernoulli probability model:

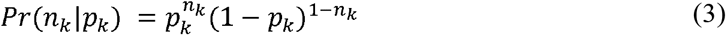

where p_k_, the probability of performing the subtask successfully at repetition k, was related to MI_k_ by a logistic function:

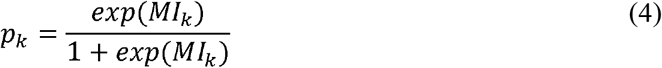

ensuring that p_k_ was constrained in [0, 1]. Furthermore, this formulation guaranteed that p_k_ would approach 1 with increasing MI.

The model parameters {α_j_, β_j_, σ_δ,j_, σ_□_, p_k_} were estimated for each individual subject using the recordings of r_j,k_ and n_k_ (i.e., kinematic recordings from the robotic device, see Section 2.4) and by applying Bayesian Monte Carlo Markov Chain methods. The estimation of the parameters resulted in an estimate for MI. In order to ensure accuracy of the model, it was necessary that the number of recordings of r_j,k_ and n_k_ exceeded the number of parameters. Based on simulations performed with varying number of data points (see *Supplementary material*), the minimum number of data points for MI estimation was set to 8. In order to validate the capability of the proposed approach to appropriately capture variable dynamics of the performance measures, we simulated different rehabilitation scenarios under varying conditions (see *Supplementary material*). As we aimed at estimating MI at subtask level, separate MI models were used for each movement direction of the training exercise.

### 5.2 Performance measures

Previous studies have shown that mechanisms of post-stroke recovery can be described by factors related to movement speed, smoothness, and efficiency [28, 47, 49]. Unlike physiological signals, these kinematic performance measures can be easily recorded and processed in real-time, promoting their use in clinical settings. In this study, we selected two continuous performance variables r_j_ for the use with the MI model: i) the mean velocity of a movement (MV) and ii) the spectral arc length (SAL), a robust and consistent measure of movement smoothness [46]. SAL is a dimensionless measure quantifying movement smoothness by negative values, where higher absolute values are related to jerkier movements. Regarding rehabilitation training, values of SAL close to zero are desirable, as well as high values of MV. Both measures were computed from the Cartesian coordinates of the three-dimensional trajectory of the robotic handle (see section *D. Robotic exoskeleton and motor task*). The discrete variable n_k_, instead, was denoted as success (SUCC) and defined separately for the experiments with the healthy participants and the patients. For the patients, the value of SUCC was determined by the robotic assistance (i.e., SUCC = 1 if the patient performed the movement without robotic assistance, SUCC = 0 otherwise). On the other hand, the healthy participants were expected not to rely on the robotic assistance, although it was also provided if necessary. This assumption was supported by preliminary experiments with healthy subjects (see *Supplementary*). Therefore, in order to have an equivalent discrete variable for the experiment with healthy subjects, we defined the value of SUCC based on the execution time (i.e., SUCC = 1 if a healthy participant completed the movement within the time threshold tth, SUCC = 0 otherwise). The time threshold tth was set to 4 seconds based on preliminary experiments with healthy subjects (see *Supplementary*).

### 5.3 Personalization routine

Using the model described in the previous section, MI was continuously tracked for each subtask (i.e., single movement towards target, see Section 5.4) and used to implement a personalized training routine. At the beginning of the training, we identified the subject-specific difficulty level for each subtask of the training exercise based on an initial assessment of the performance measures. The subtasks were then ordered by increasing difficulty and the easiest ones were selected as the initial training set (see section 5.6 for details on the ordering of the single movements). During the training, a subtask was removed from the set of current training subtasks when the MI estimates for this movement exceeded a given threshold and approached a plateau. Specifically, the probability of performing the subtask successfully (p_k_, see Section 5.1) had to be greater than 0.5, and the difference between two consecutive MI values (i.e., between two repetitions of the same subtask) had to be smaller than 5% for at least four repetitions. Given the observation equation for p_k_, the former condition (p_k_> 0.5) can be equally expressed in terms of the motor improvement: MI_k_ > 0. Once these conditions were satisfied, the subtask was replaced by a more difficult one from the training queue. The removed subtask was placed back into the training queue, so that it could be reintroduced at a later stage.

### 5.4 Robotic exoskeleton and motor task

We implemented the motor improvement model and the personalization routine in the robotic upper-limb exoskeleton ALEx [39, 40]. During the experiments, the patient and the healthy participants were instructed to perform point-to-point reaching movements at their comfortable speed (Fig. 1a). All reaching movements started from the center of the workspace and the goal was to reach one of the eighteen targets distributed over a sphere of 19 cm of radius (Fig. 1b). Each movement towards a target represented a subtask. This design allowed exploiting an extensive three-dimensional workspace and provided means to easily identify all subtasks of the exercise. The sphere was positioned so that its center was aligned with the acromion of the right arm mid-way between the center of the target panel and the subject’s acromion. The targets were displayed on a screen mounted in front of the subjects and visual feedback was provided by means of a cursor mapping the position of the exoskeleton’s handle to the screen. In order to preserve the depth perception, the dimensions of the target spheres were modified in accordance with their position in the 3D space. If a subject was unable to reach a target (i.e., the subject did not move for more than 3 seconds), ALEx activated its assistance mode to guide the subject towards the target according to a minimum jerk speed profile [62].

### 5.5 Participants

#### 5.5.1 Healthy participants

Seventeen right-handed subjects (eight males, nine females, 25.4 ± 3.3 years old) participated in the experimental validation of our approach. The participants did not present any evidence or known history of skeletal and neurological diseases and they exhibited normal ranges of motion and muscle strength. All participants gave their informed consent to participate in the study, which had been previously approved by the Commission Cantonale d’Éthique de la Recherche Genève (CCER, Geneva, Switzerland, 2017-00504).

#### 5.5.2 Subacute stroke patients

Two subacute stroke patients from the inpatient unit of the Hôpitaux Universitaires de Genève (HUG, Geneva, Switzerland) were included in the study. A summary of the patient information is reported in Table 1. Both patients suffered from a right hemiplegia with at least 10° of residual motion in shoulder and elbow joints. The patients were enrolled in the study within two to eight weeks after the stroke and underwent a therapy following the adaptive robotic rehabilitation protocol described in section 5.6. The patients received the robot-aided treatment in addition to a standard non-robotic rehabilitation therapy: each patient received two sessions of 30 minutes of physical therapy per day, five days per week, as well as five sessions of 30 minutes of occupational therapy per week, on an inpatient basis, for 8 to 16 weeks. This was followed by an outpatient treatment of 1-4 hours of physical and occupational therapy per week. All patients gave their informed consent to participate in the study. This study is registered in ClinicalTrials.gov (NCT02770300) and the experimental protocols were approved by Swissmedic and Swissethics.

**Table 1.**
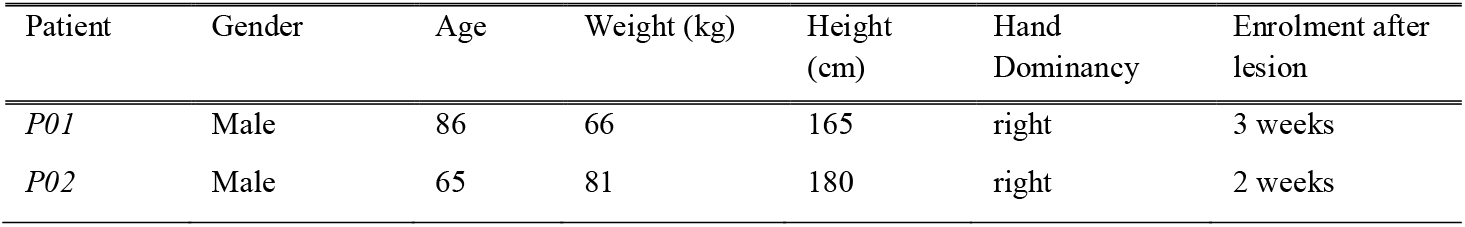
Demographics and information of the stroke patients included in the study

| Patient | Gender | Age | Weight (kg) | Height (cm) | Hand Dominancy | Enrolment after lesion |
| --- | --- | --- | --- | --- | --- | --- |
| *P01* | Male | 86 | 66 | 165 | right | 3 weeks |
| *P02* | Male | 65 | 81 | 180 | right | 2 weeks |

### 5.6 Experimental protocols

#### 5.6.1 Healthy participants

The healthy participants attended a single experimental session, which comprised seven blocks of reaching movements (Fig. 1c). Breaks were allowed between the blocks to prevent fatigue. The session started with an initial assessment block consisting of three runs (A_I,1-3_). During each run all 18 targets were presented once and in a randomized order. The purpose of the assessment block was i) to allow familiarization with the robotic system and the motor task and ii) to record a baseline for the performance measures. This block was followed by five blocks B_1-5_ during which the visual feedback was inverted (i.e., an upward movement was displayed as downward and vice versa, likewise for left/right and forward/backward movements). This visual manipulation was introduced to induce motor performances with temporal dynamics resembling the ones observed in robot-aided rehabilitation of stroke patients [28, 37, 38]. At the onset of the five inversion blocks, participants were not informed about the manipulation of the visual feedback, but they were told that the task difficulty was changed. Each of the five inversion blocks B_1-5_ consisted of five runs, each one composed of eight point-to-point reaching movements for a total of 40 reaching movements per block.

The initial set of training targets for each participant was generated following a semi-randomized procedure: it always contained the six on-axis targets (i.e., targets 1, 3, 5, 7, 10 and 13, see Fig. 1b) and two randomly selected off-axis targets (i.e., targets 2, 4, 6, 8, 11, 14, 15, 16, 17 and 18). The presentation order of the eight initial training targets was randomized. The remaining ten off-axis targets were placed randomly in the training queue. Previous studies using planar setups [33, 63] demonstrated that participants showed better performance for targets lying on the axis perpendicular to the inversion. Although in this study we used a three-dimensional setup, we also hypothesized that participants would have less difficulty with the on-axis targets, as they involved inversions in only one dimension (in contrast to inversions in two dimensions for the off-axis targets).

During the five inversion blocks B_1-5_, a target was removed from the current set of training targets if the MI estimates for this subtask satisfied the replacement conditions (see section 5.3). In this case, the target was replaced by the next one in the training queue. The inversion blocks B_1-5_ were followed by a final assessment block which was composed of three runs (A_F,1-3_) and followed the same procedure as the initial assessment block (i.e., neither visual manipulation nor personalization were applied). The data acquired during the assessment blocks (i.e., A_I,1-3_ and A_F,1-3_) were not considered for the MI estimation.

#### 5.6.2 Subacute stroke patients

The experimental protocol for the patients consisted of four weeks of robot-aided rehabilitation therapy (Fig. 1d), with three sessions of 30 minutes per week. The training comprised the regular point-to-point reaching task (see section 5.4). In order to evaluate the outcome of their rehabilitation training, the patients completed two assessment sessions before (A_I,1-2_) and after (A_F,1-2_) the therapy. The initial assessment sessions A_I,1-2_ were completed two weeks and one week before the beginning of the therapy. The final assessment sessions A_F,1-2_ were completed one week and one month after the end of the therapy. During the initial and final assessment sessions, all eighteen targets of the point-to-point reaching task were presented to the patients in a randomized order. The total amount of reaching movements for each session was determined by the physical therapist while encouraging the patient to perform as many movements as possible. In addition, the patients were evaluated using the upper extremity section of the Fugl-Meyer assessment (FMA-UE, [64]).

For the treatment sessions, we first identified the patient-specific difficulty for each of the 18 targets following the initial assessment sessions A_I,1-2_. Specifically, we analyzed the mean values of MV, SAL and SUCC for each of the eighteen training targets. The targets were first ordered by descending (i.e., starting from easier targets) mean values of SUCC (rate of SUCC). If several targets had equal values for the rate of SUCC, the order amongst them was determined by their mean values for MV and SAL, while giving both measures equal weight. The first eight targets of the resulting list were selected as the initial training targets. The remaining targets were placed in a training queue while conserving the determined order of difficulty. During the therapy (W_1_-W_4_, see Fig. 1), MI was continuously estimated for each training target separately. The replacement of a training target based on the MI estimates followed the procedure presented in Section 5.3. The current set of training targets was saved after the completion of each training session, ensuring continuity between sessions. The total amount of reaching movements for each session was determined by the physical therapist while encouraging the patient to perform as many movements as possible.

### 5.7 Statistical analysis

A two-sample t-test was used to compare the performance differences between two groups within the healthy population (fast and slow adapters). A two-way ANOVA was used to assess the interaction effects of visual manipulation (introduced between A_3_ and B_1_) and adaptation speed (fast and slow adapters) in healthy participants. A paired t-test was used to compare the performances between different time points for the patients performing the pilot test. A significance level of 0.05 was used for all analyses. All analyses were performed using MATLAB (The MathWorks, Natick, Massachusetts).

## List of abbreviations

MI: Motor improvement
MV: Movement velocity
SAL: Spectral arc length
SUCC: Success
FMA-UE: Fugl-Meyer assessment for upper extremities

## Declarations

### Ethics approval and consent to participate

All participants gave their informed consent to participate in the study, which had been previously approved by the Commission Cantonale d’Éthique de la Recherche Genève (CCER, Geneva, Switzerland, 2017-00504).

### Consent for publication

All participants signed an informed consent to the use of all coded data collected during the study in scientific publications.

### Availability of data and material

Since the data used in this study includes data collected in a clinical trial with patients, the data will not be shared.

### Authors’ contributions

C.G. designed the model, carried out experiments, analysed data and wrote the paper;

E.P. designed the model, carried out experiments, analysed data and wrote the paper;

N.K. designed the model, carried out experiments, analysed data and wrote the paper;

C.P. designed the model, carried out experiments, analysed data and wrote the paper;

A.P. designed the model and wrote the paper;

M.C. carried out experiments and wrote the paper;

J.M. carried out experiments and wrote the paper;

C.M. carried out experiments and wrote the paper;

P.N. carried out experiments and wrote the paper;

A.G. designed the model and wrote the paper;

S.M. designed the model and wrote the paper;

## Acknowledgements

The authors would like to thank all the volunteers and the patients enrolled in the study. We would also like to thank Wearable Robotics and PERCRO for their support and expertise.

---

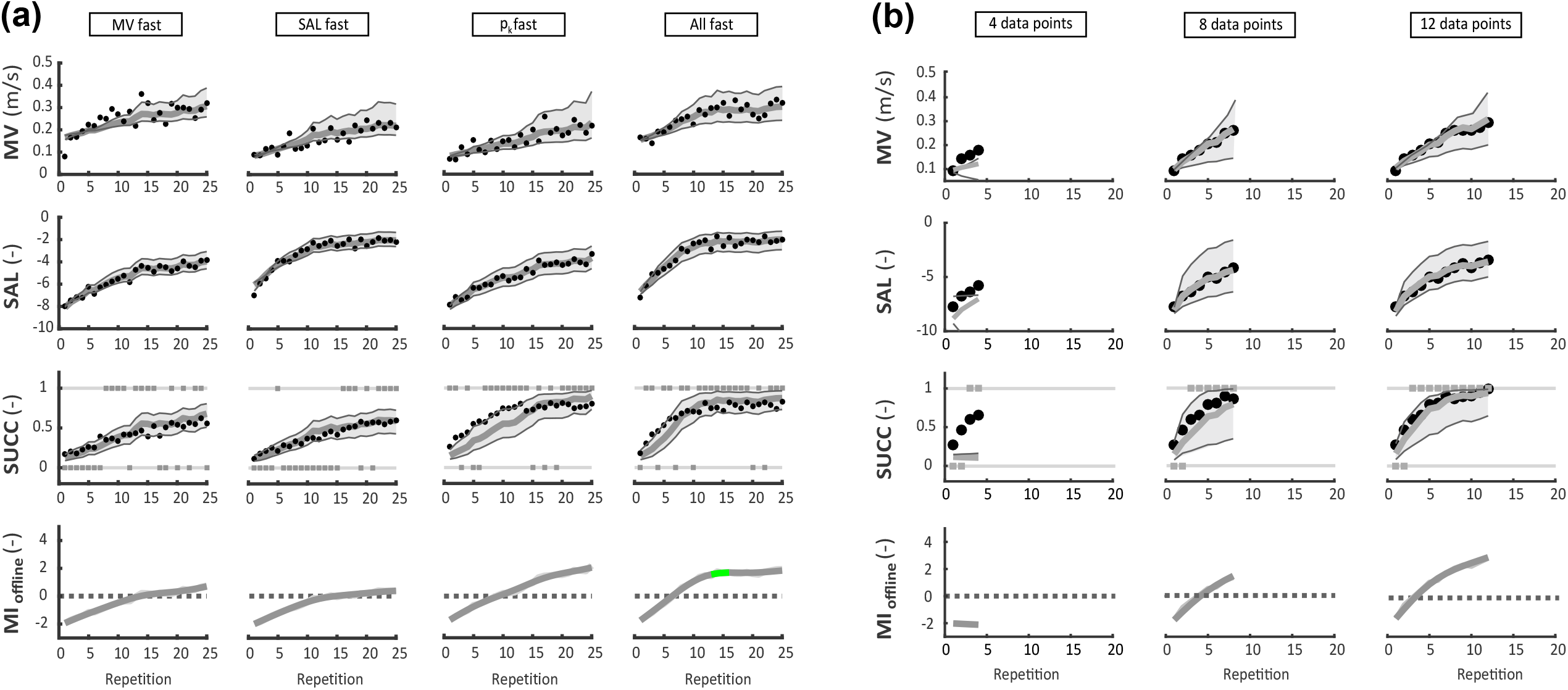

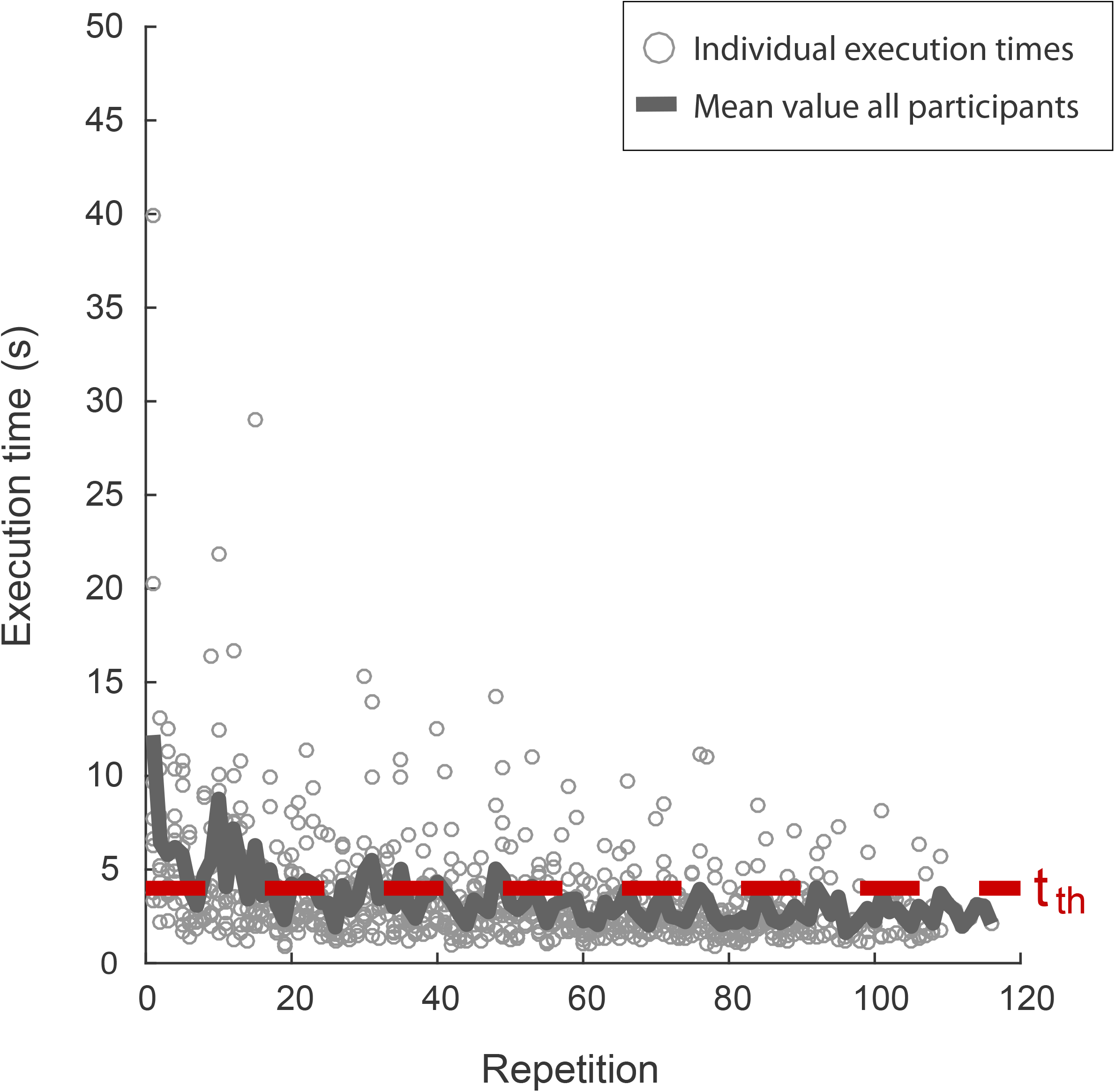

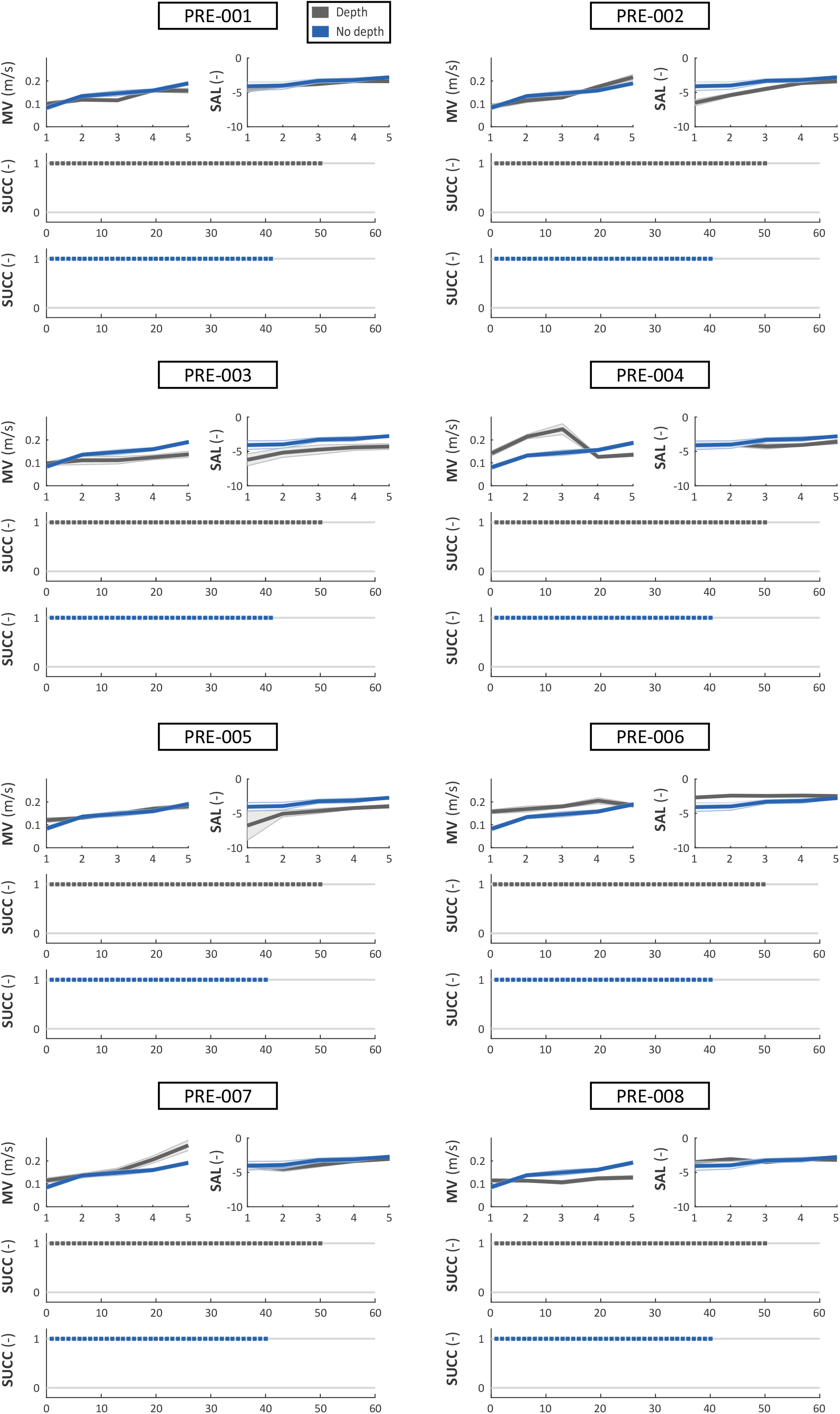

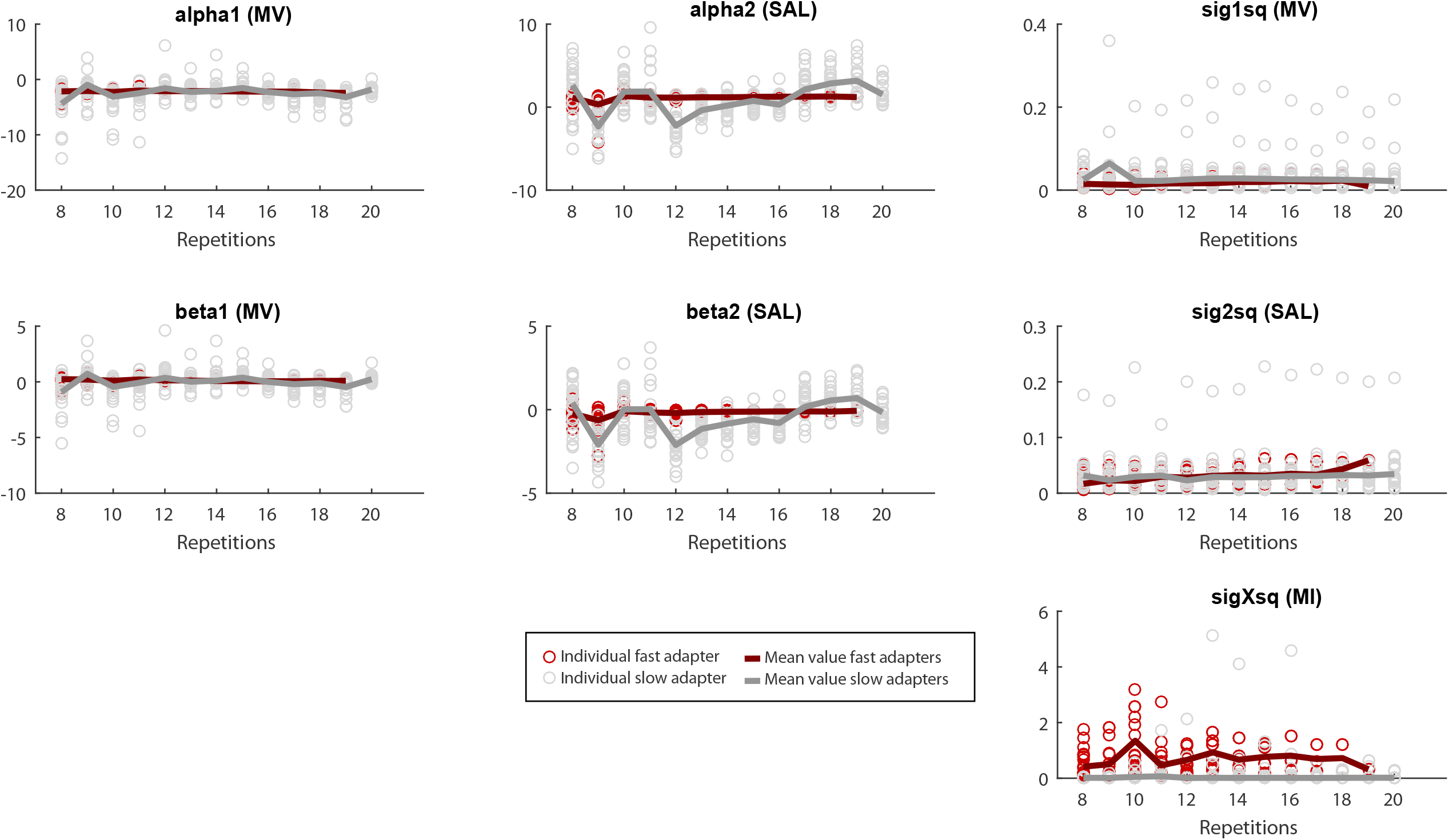

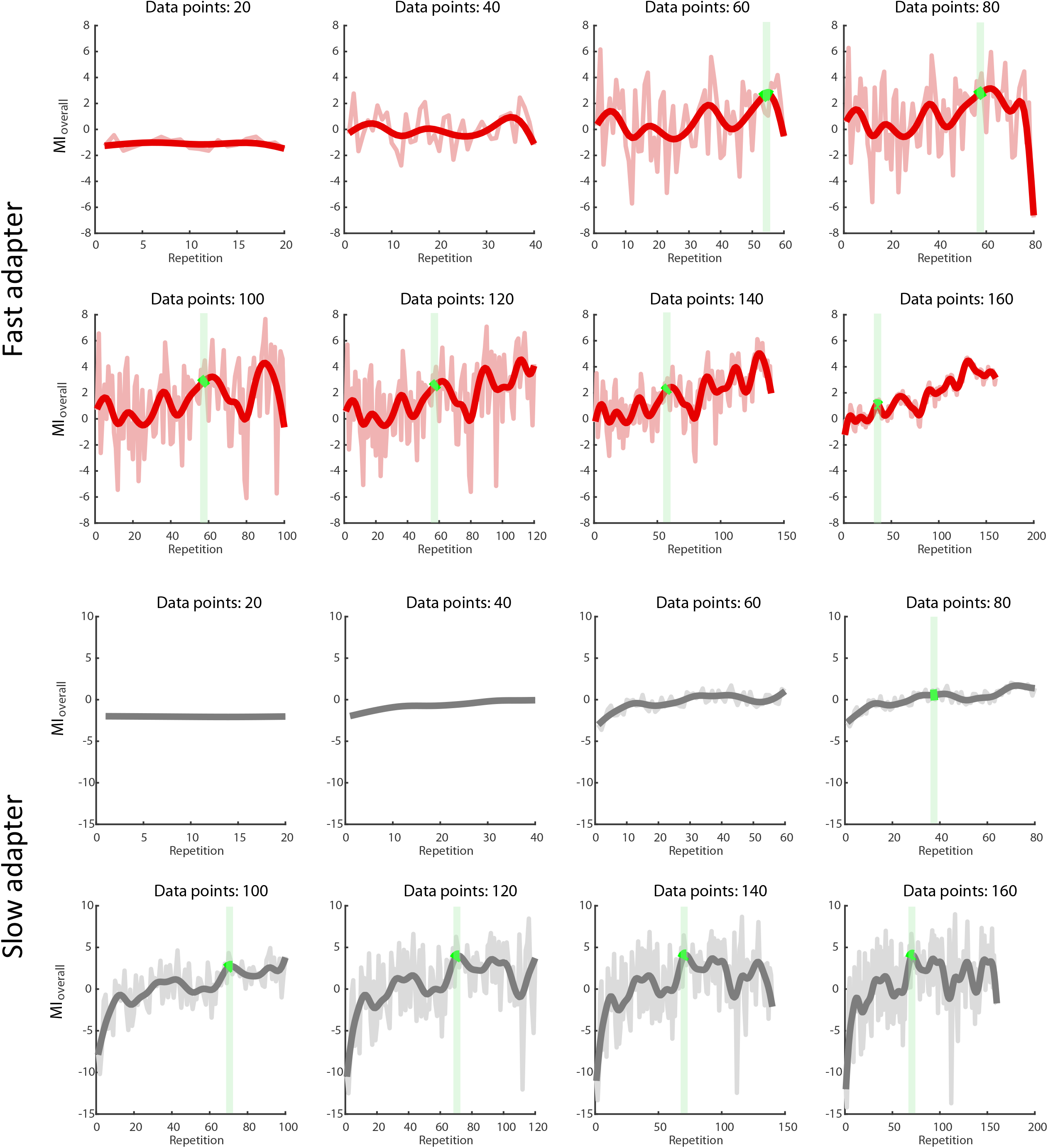

