## Supplementary for "Motor improvement estimation and task adaptation for personalized robot-aided therapy: a feasibility study"

**Additional analysis of performance measures**

In order to demonstrate the capability of the proposed model to capture varying performance measures dynamics, we simulated different rehabilitation scenarios under varying conditions (Fig. S1a). Specifically, data were generated for the three variables MV, SAL, and $p_{k}$ using an exponential equation:

$$r_{j,k} = r_{j,end}-r_{j,start} exp(-\frac{k}{\tau_{j}})$$

(1)

where $j = 1,2,3$ are the different performance measures and $k = 1,...,25$ are the different repetitions of a movement. $r_{j,end}$ and $r_{j,start}$ are parameters used to set the desired initial and final values of each performance measure.$\tau_{j}$ is the individual time constant for each performance measure. The equation was used to simulate the data of MV, SAL, and $p_{k}$ for 25 repetitions of the movement towards the same target. The values for SUCC were deduced by using the values of $p_{k}$ and a Bernoulli distribution model. We ran the simulations under four conditions: in the first three conditions, the time constant of one performance measure was reduced to$\tau= 5$, while the other two were kept at $\tau= 15$. In the fourth condition, the time constants for all three measures were reduced to $\tau= 5$. For all conditions, we obtained approximations of the simulated data by inserting the estimates of the unknown model parameters into the observation equations. Moreover, we calculated the 95% confidence intervals of the approximations and the corresponding motor improvement estimates. The results of the simulations illustrate the capability of the proposed model to capture varying dynamics of the performance measures properly. The simulated data lie within the 95% confidence intervals of the approximations for the most part. Moreover, the only condition where the requirement for a replacement is met is the one where all performance measures are simulated with low time constants and quickly reach a plateau, highlighting the fact that a replacement is only suggested by the algorithm when no further improvement is expected.

Moreover, we simulated MI estimation for lower number of data points (Fig. S1b). The simulations presented in Fig. S1b were run with varying amount of data (4, 8 and 12 data points) from the same data set. We observed that the simulated data were mostly in the 95% confidence interval of the model estimates for MV, SAL and p_k_ when 8 or 12 data points were used for the estimation. However, this was not the case when only 4 data points were considered for calculating the estimates. Based on this analysis, we have set the minimum number of data points necessary for MI estimation to 8.

**Figure S1**. Simulated data for motor improvement estimation under varying conditions. (a) Data shows the simulated performance measures and the corresponding motor improvement estimates under four different conditions for 25 repetitions of the same movement. In the first column, the time constant for the mean velocity (MV) was reduced to $\tau=5$ repetitions ($\tau=15$ repetitions for the remaining two measures), in the second column the time constant of the spectral arch length (SAL) was reduced to $\tau=5$ repetitions and in the third column the time constant reduced to $\tau=5$ repetitions was the one of the probability of success (p_k_). In the last column, the time constant for all performance measures was set to $\tau=5$ repetitions. The first two rows show simulated data (black dots) for MV and SAL. The third row depicts the simulated data for p_k_ (black dots) and the corresponding discrete performance measures SUCC (grey squares) deduced from p_k_ using a Bernoulli distribution model. Grey lines show approximations of the performance measures using the estimated parameters resulting from the algorithm. Shaded area depicts 95% confidence interval of the approximations. The last row shows the resulting offline motor improvement estimates (MI_offline_) using an offline implementation of the model. Dotted lines depict necessary condition (MI_offline_ > 0) for target replacement. Green area indicates the time span where the algorithm detects a performance plateau and suggests a target replacement. (b) Simulated motor improvement estimates provided by the model based on 4, 8 and 12 data points of the same data set. The first two rows show simulated data (black dots) for MV and SAL. The third row depicts the simulated data for p_k_ (black dots) and the corresponding discrete performance measures SUCC (grey squares) deduced from p_k_ using a Bernoulli distribution model. Grey lines show approximations of the performance measures using the estimated parameters resulting from the algorithm. Shaded area depicts 95% confidence interval of the approximations. The last row shows the resulting offline motor improvement estimates (MI_offline_) using an offline implementation of the model. Dotted lines depict necessary condition (MI_offline_ > 0) for target replacement.

**Preliminary experiments with healthy participants**

**Figure S2**. Execution time of eight healthy participants (seven females, a male, 54.8 ± 13.8 years old) performing the regular point-to-point reaching task. Data for each participant is pooled for all movement directions (i.e., for all targets) and presented in chronological order (grey circles). Solid grey line shows evolution of execution time averaged over all eight participants. Red dashed line depicts the time threshold (t_th_ = 4s) used to determine the discrete performance measure SUCC in the visually manipulated reaching task. t_th_ was selected as the upper bound for the average execution time after the 50th repetition.

**Figure S3.** Performance measures of eight healthy participants (seven females, a male, 54.8 ± 13.8 years old) performing the regular point-to-point reaching task (same data reported in Fig. S2). First row shows the mean values and standard error of the mean of MV and SAL for five repetitions of no-depth targets (blue, targets 1-8) and depth targets (grey, targets 9-18). Data for each repetition is averaged for all targets of a class. Second and third row show the data of SUCC for no-depth (blue) and depth targets (grey). The data for each class of targets is presented chronologically for each repetition of the movements. SUCC was defined by the usage of the robot assistance (i.e., SUCC = 1 if the participant performed the movement without robotic assistance, SUCC = 0 otherwise). Overall, performances were not different for depth and no-depth targets, confirming that the participants could properly perceive the depth.

**Additional analysis of model parameters**

We have performed additional analyses to illustrate the temporal dynamics of the model parameters for both fast and slow adapters (Fig. S4). With increasing number of repetitions, all model parameters converged towards final values, indicating that the recorded data fit well the model. Interestingly, the model parameters for the fast adapters seemed to converge faster, probably because in this group participants showed faster improvements in the performance measures, which would fit the chosen observation models. When looking at the final mean values of the model parameters for both groups we did not observe notable differences, indicating that on average, motor improvement models were similar for both groups and that improvement could be observed only in changes of the MI estimates. The only remarkable difference was found for σϵ which appeared to be higher for the fast adapters, reflecting the improvements in MI values for this group. Finally, values for σ_δ,j_ remained bounded in a range of reasonable values, indicating a limited influence of the gaussian noise terms δ_j_ on the motor improvement estimates.

**Figure S4.** Model parameters estimated for participants doing the visually inverted reaching task (n=17). Paramters were estimated using Baysian Monte Carlo Markov Chain methods with three independent estimation chains and different initial guesses. Data is presented for movements towards all targets for all subjects of the fast adapter (red circles) and slow adapter (grey cirlces) groups. Red and grey lines indicate average parameter values for both groups. Parameters were calculated for each number of repetitions towards a target (minimum 8 repetitions, maximum 20 repetitions).

Moreover, we calculated overall MI estimates for the two subjects presented as examples in Fig. 3. Therefore, we merged the data from movements towards all training targets and calculated the MI estimates based on these combined data sets. Although an increase in MI can be observed in both subjects, we observed that the overall MI estimates for both subjects appeared to be noisier and less descriptive, since now data of movements towards easy and difficult targets are pooled together. Indeed, basing the analysis on the overall MI estimates, the recoveries of the movements presented in Fig. 3, are obscured by the inferior performances recorded for the difficult targets. Moreover, the detection of performance plateaus would not correspond to the actual performances for each subtask. As a result, some subtasks would be kept too long, while others would be replaced too early, potentially leading to a less efficient training schedule. For instance, the overall MI estimate for the slow adapter suggest a performance plateau already after 39 reptitions (corresponds to approximately 5 repetitions for each subtask). However, when looking at the performance measures of this subject for target 13 seperately, it is clear that a replacement of this target after 5 repetitions would have been too soon. We therefore believe that this analysis further supports our approach to specifically consider MI estimation at subtask level.

**Figure S5.** Overall MI estimates for fast (red) and slow adapter (grey) presented as examples in Fig. 3. Overall MI estimated were based on data chronologically merged from all training targets and were calculated for the first 20, 40, 60, 80, 100, 120, 140 and 160 data points. Data for MI estimates were low-pass filtered for visualization purposes (raw data shown in light red/grey). Green area indicates the time span where the algorithm detects a performance plateau and suggests a target replacement.
